# State-Space Compression Enables Desktop-Scale Hit Discovery for Intrinsically Disordered α-Synuclein

**DOI:** 10.64898/2026.06.22.733879

**Authors:** Dae Hoon Kim, Gerelt-Od Khenmedekh, Jihyeon Park, Sangjune Kim

## Abstract

The accessible chemical space dwarfs any tractable screening budget, and most artificial intelligence drug discovery pipelines respond by docking and ranking a small sublibrary. The resulting hit list is agnostic to selectivity, brain penetration, toxicity, synthetic accessibility, and chemical novelty. We present ISTP-DPISO DrugEngine, an end-to-end engine developed by ISTP Tech that integrates the Local Information Criticality Principle (LICP) with a Discrete Phase-Interference Search Operator (DPISO). We demonstrate the engine on the intrinsically disordered protein (IDP) α-synuclein, whose non-amyloid-component (NAC, residues 61–95) drives Parkinson-associated aggregation. The resulting LICP active set focuses the expensive LICP-DPISO scoring: in a production-scale run, the engine compressed a ~8.46×10^8^-molecule mirror to a 10,000,000-molecule active set (~85-fold) before scoring, then converged to a compact, safety-gated shortlist plus de novo designs. The entire campaign ran on a single desktop workstation, without any high-performance-computing cluster. Three engine-prioritized, commercially available candidates (2-D08, Uralenol, Herbacetin) and an (−)-epigallocatechin gallate (EGCG) positive control were then tested in a thioflavin-T (ThT) aggregation assay at 100 µM: all three engine-nominated candidates suppressed α-synuclein aggregation, giving perfect prospective inhibitor-call concordance (3/3 nominated); together with the EGCG positive control, all four assayed compounds inhibited aggregation (4/4 total), two by ≥80% plateau reduction. ISTP-DPISO DrugEngine reframes virtual screening from post-hoc score fusion to a single, state-space-compressed, safety-gated, experimentally validated discovery pipeline.

## Introduction

Conventional virtual screening is limited by the combinatorial explosion of the accessible chemical space, and the drug-like chemical universe is estimated at ~10^60^ molecules, whereas even ultra-large make-on-demand libraries (10^9^–10^10^) and practical docking campaigns (10^5^–10^7^ evaluated) sample only a vanishing fraction^1^. The dominant paradigm couples a target structure to a docking engine and ranks a tractable sub-library by predicted binding affinity^2–6^. Recent reviews have surveyed the rapid expansion of artificial intelligence methods across early discovery^7^. Refinements such as consensus scoring and score fusion improve enrichment; however, they share a structural limitation in that the pipeline ends at a ranked hit. Selectivity against off-targets, brain penetration, metabolic and toxicological liability, synthetic accessibility, and critically, whether a proposed molecule is chemically novel or merely a rediscovery of known chemotypes, are all deferred to manual downstream triage. For intrinsically disordered proteins (IDPs), which lack a stable pocket and populate a conformational ensemble, the docking-centric paradigm is weaker still^8^.

α-Synuclein is the prototypical disease-relevant IDP: its aggregation into amyloid fibrils underlies Parkinson’s disease and related synucleinopathies^9,10^, and its non-amyloid-component (NAC, residues 61–95) is the minimal aggregation-driving segment^11^. Several natural products—EGCG ((−)-epigallocatechin gallate)^12,13^, baicalein^14^ and related polyphenols—remodel α-synuclein aggregation, and broader small-molecule screens have identified additional assembly inhibitors^15^, but a prospective, physics-grounded engine that designs, filters and prioritizes such inhibitors end-to-end has been lacking.

Here, we describe the ISTP-DPISO DrugEngine, organized as a single lineage—the ISTP interpretive framework, the Local Information Criticality Principle (LICP) compression criterion, and the Discrete Phase-Interference Search Operator (DPISO)—derived from our prior state-space compression studies^16,17^. LICP compresses the candidate state space into an information-bearing active set. DPISO then searches this active set using phase accumulation, constructive amplification, and destructive pruning. The result is a single eight-stage pipeline that produces a wet-lab-ready, safety-gated, commercially sourced, novelty-annotated shortlist. The engine is the contribution; α-synuclein is the validation case. The three nominated commercial candidates inhibited α-synuclein aggregation in a thioflavin-T (ThT) assay, with EGCG behaving as the expected positive control, closing the loop from a physics-based design to experimental confirmation. Conceptually, most pipelines reduce the search space implicitly through library design choices made before any score is computed; the ISTP-DPISO DrugEngine explicitly reduces it as a measured, reported compression step inside the pipeline.

## Results

### An eight-stage physics-constrained discovery engine

ISTP-DPISO DrugEngine replaces the conventional target→docking→hit chain with eight sequential, physically motivated stages (Fig. 1). Stage 1 defines target hotspots via the Protein Genesis residue-graph module (for α-synuclein, the NAC β-sheet core V66–V77, the G68–A69 nucleation hinge and the E83 salt-bridge anchor). Stage 2 performs a DPISO search across each candidate’s compressed conformational manifold by phase interference, whereas Stage 3 applies the LICP stability gate, retaining only candidates whose interaction manifold is information-theoretically stable. Stage 4 involves the growth of new chemotypes by de novo fragment assembly with scaffold dead-branch termination^3^. Stages 5–7 are the safety and developability gates: cross-reactivity rejection against a 242-hit-point anti-target panel, blood-brain barrier (BBB) and absorption, metabolism, excretion, and toxicity (ADMET) filtering, and commercial availability annotation. Stage 8 runs a ZINC reverse novelty search to label each candidate as a novel chemotype or a known analog. The engine is configuration driven; the engine code is never modified between targets (only the JSON target definition changes), which underwrites the reproducibility.

**Figure 1.**
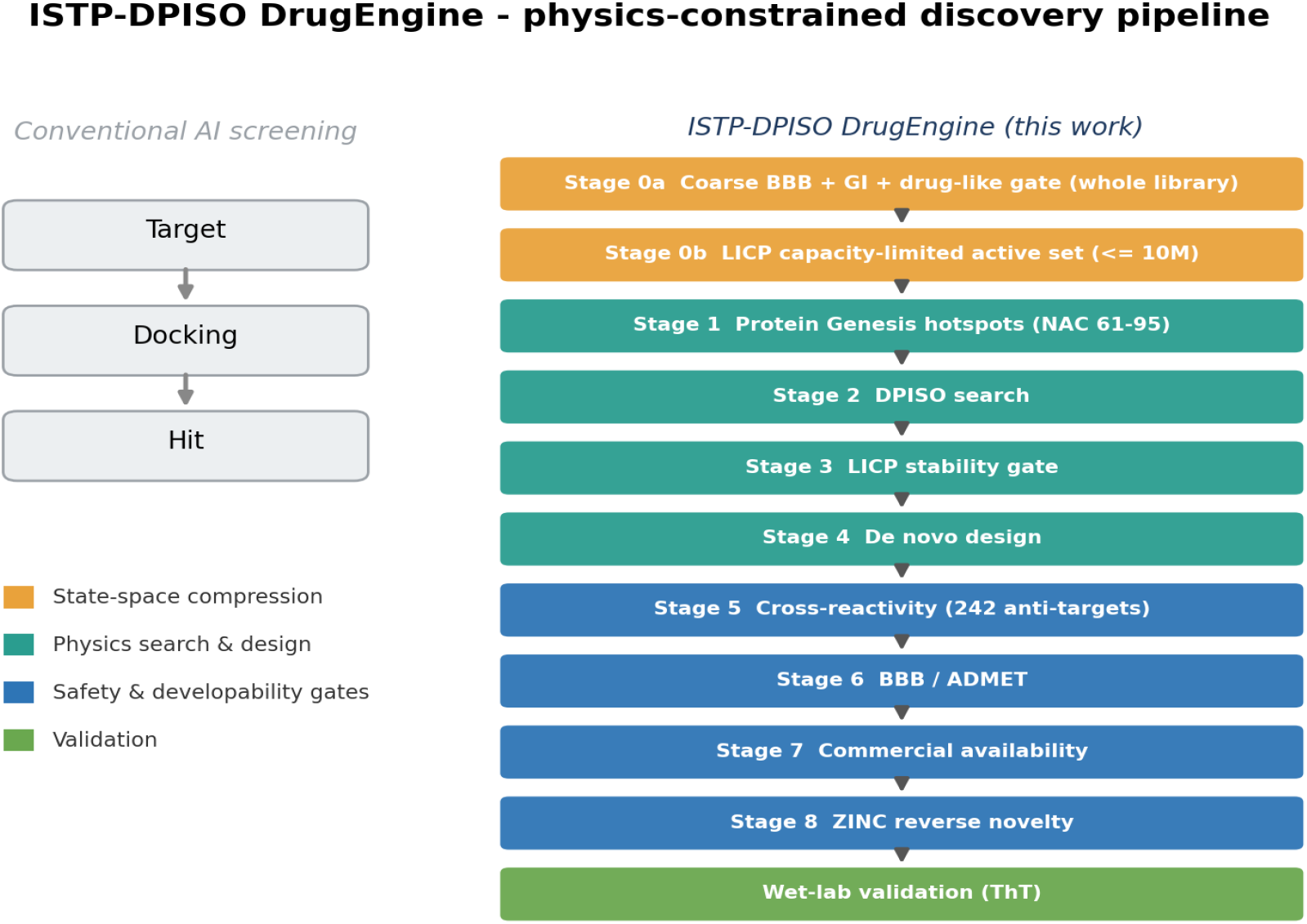
Engine architecture. Conventional AI screening (left) terminates at a ranked hit. ISTP-DPISO DrugEngine (right) threads each candidate through eight physically motivated stages—Genesis hotspot, DPISO search, LICP stability, de novo generation, cross-reactivity rejection (242 anti-targets), BBB/ADMET, commercial availability, and ZINC reverse novelty—ending in wet-lab validation.

**Figure 2.**
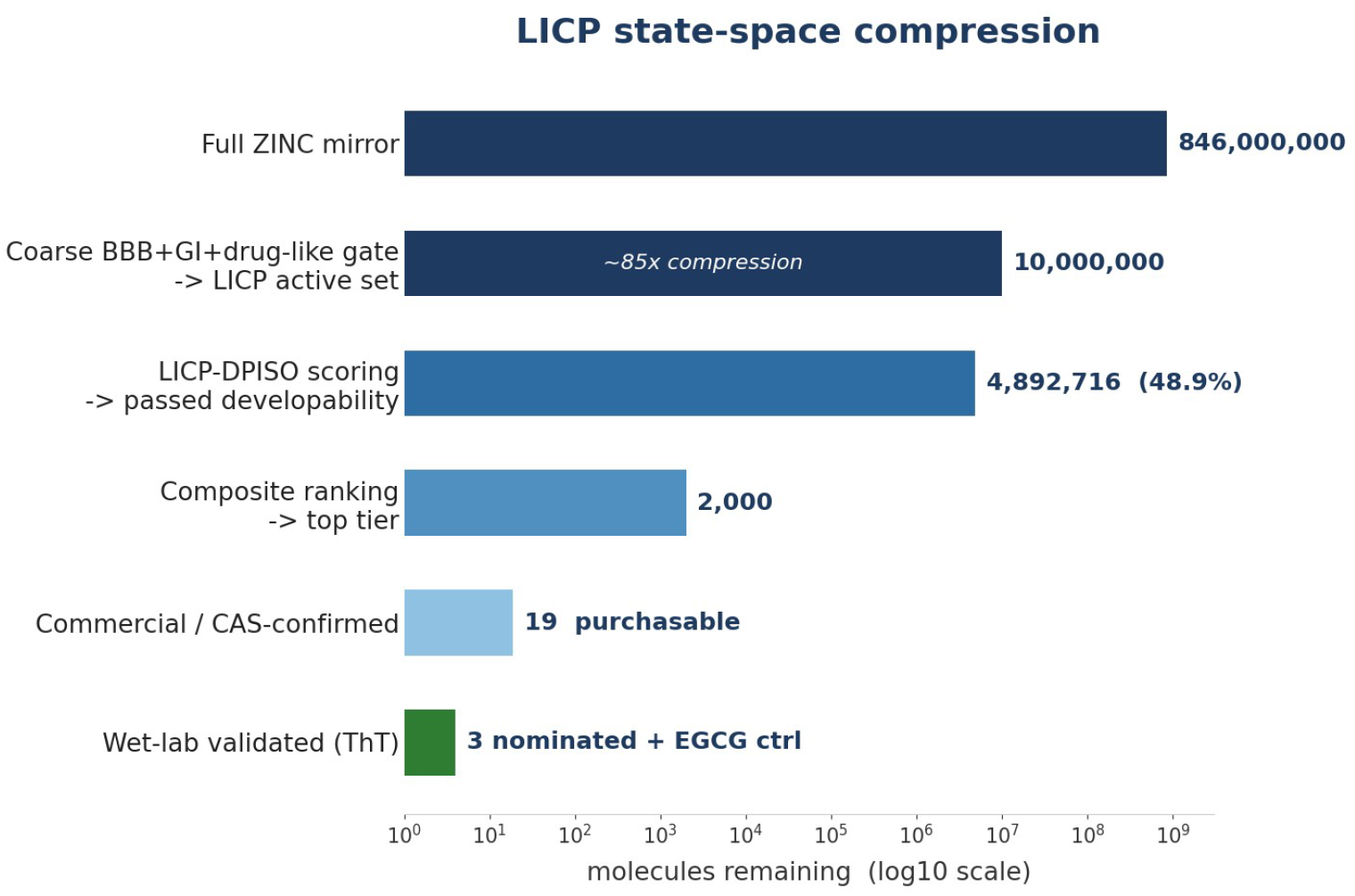
LICP state-space compression. The full ~8.46×10^8^-molecule ZINC mirror is reduced explicitly at each stage: a cheap BBB+GI+drug-like coarse gate yields a 10,000,000-molecule LICP active set (~85×); LICP-DPISO scoring passes 4,892,716 (48.9%); composite ranking and the safety gates narrow this to a top tier, 19 purchasable CAS-confirmed compounds, and 3 engine-nominated candidates plus an EGCG positive control assayed by ThT. The decisive lever is the explicit compression step, not a faster scorer. Log_10_

### The ISTP composite scoring system

Each candidate is ranked by a single composite score that fuses engine physics with developability. In IDP mode (used for alpha-synuclein) the score is 0.30 x Genesis-landscape + 0.20 x persistence + 0.20 x DPISO phase-interference + 0.15 x LICP stability + 0.10 x quantitative estimate of drug-likeness (QED) + 0.05 x synthetic accessibility, reduced by an ADMET penalty (0.03 per soft toxicity flag, capped at 0.15) and multiplied by a hard-filter gate. The target-aware score combines this with pocket complementarity as target_combined = 0.72 × composite + 0.28 × pocket-fit, where pocket-fit is the small-molecule match to the Genesis-defined NAC hotspots. Analogous weightings (emphasizing hotspot contact and interface retention) define the orthosteric and protein-protein interaction modes.

Selectivity is scored explicitly: for every candidate the engine computes a selectivity gap (target pocket-fit minus the maximum off-target pocket-fit across the 242-hit-point anti-target panel) and assigns a LOW/MODERATE/HIGH cross-reactivity label on the same scale -- target pocket-fit is never compared against an off-target combined score. The composite score was validated by calibration before any screening; against alpha-synuclein, active/decoy discrimination reached an AUC of 0.955 (Cohen’s d 2.62; 95% bootstrap confidence interval reported in the SI), and three literature inhibitors were independently re-ranked near the top of a 283,383-compound blind screen (luteolin rank 11, quercetin rank 79, and baicalein rank 105), confirming that the score tracks genuine anti-aggregation activity.

### Calibration and the screening funnel

Applied to the ZINC15 In-Stock Reactive library (283,383 compounds), the funnel passed 200,497 molecules through the Stage-0 developability gate, ranked the survivors by the composite score and retained the top 2,000 (LICP-DPISO). Nineteen Chemical Abstracts Service (CAS)-confirmed, purchasable chemotypes were identified within the top 200, and an eleven-compound validation panel -- three independently-recovered known actives, five engine-nominated, α-synuclein-unreported commercial compounds and controls -- was advanced to wet-lab assay (Supplementary Fig. 1). For ultra-large libraries (the local ZINC mirror holds ~ 8.5 × 108 molecules across 1,313 tranches), the same composite score is reached after a cheap BBB + gastrointestinal absorption gate populates an LICP capacity-limited active set (production ceiling 10,000,000), so the expensive LICP-DPISO scoring never touches the full set. In a production-scale run on this principle, the coarse gate compressed the ~8.46×10^8^-molecule ZINC mirror into a 10,000,000-molecule LICP active set; DPISO scoring of that set passed 4,892,716 molecules (48.9%) through the engine developability filter (Supplementary Table 4).

### Safety-aware multi-stage filtering

A defining feature of the engine is that selectivity and developability are gates inside the pipeline and not afterthoughts. In the cross-reactivity stage, each surviving candidate was screened against a panel of 242 anti-target hit points derived from 100 seed proteins (kinase ATP pockets, GPCRs, nuclear receptors, and other liability targets). For an IDP target, this stage is deliberately stringent because the NAC groove is shallow and non-specific; every one of the 200 top-ranked candidates (200/200, 100%) was flagged as HIGH cross-reactivity risk (mean maximum off-target pocket-fit 0.82), a biologically correct outcome for a shallow IDP groove, and the signal that motivates a de novo first strategy to escape promiscuous chemotypes. A HIGH flag denotes the predicted non-selective pocket engagement, not toxicity, and these compounds are tox-CLEAN, reflecting the shallow, non-specific NAC groove intrinsic to IDPs. The coarse gate enforces BBB permeability as a screening filter, but the BBB call is compound-specific: a stricter BBB-tight re-scoring (preset bbb_cns) retained 535 active-set molecules as central nervous system (CNS)-developable, whereas the commercially sourced validation panel was chosen for ThT assay ability rather than CNS candidacy — among the tested compounds only 2-D08 is BBB-positive (Table 1). The commercial arm screens the in-stock subset so that survivors can be immediately ordered (Supplementary Fig. 2).

**Table 1.**
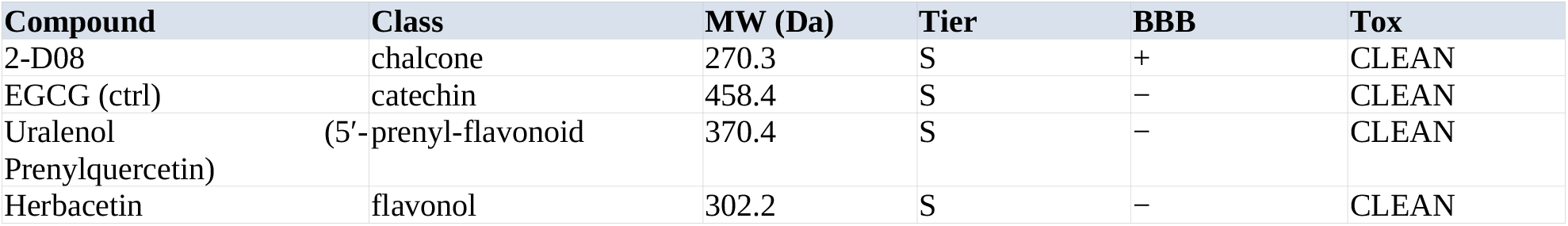
Engine-nominated Tier-S compounds advanced to wet-lab (commercial reagents).

### De novo design and novelty handling

Stage 4 grows new chemotypes from fragment seeds using directional fragment assembly with a dead-branch termination rule borrowed from LICP maze-solving; a growth branch is abandoned as soon as a child fragment scores below 80% of its parent, and scaffold families represented by a single analog are pruned (scaffold leaf-peeling). Each de novo candidate is additionally checked for novelty against the in-stock ZINC set. Genuinely new chemotypes are routed to synthesis, whereas close analogs of purchasable compounds are flagged for immediate ordering. In this run the de novo stage produced 10 peptide-binder designs, all novel by sequence, and the reverse-novelty search labelled 20 of 50 scaffold-diverse small-molecule hits as novel chemotypes (maximum Tanimoto < 0.40 against 200,000 ZINC references); full novelty values are tabulated in the SI.

### Wet-lab validation: 3/3 prospective candidates plus EGCG control

To test the engine prospectively, three engine-prioritized commercial candidates—2-D08 (chalcone), Uralenol (prenyl-flavonoid) and Herbacetin (flavonol)—together with an EGCG (catechin) positive control were assayed for their effect on α-synuclein aggregation by thioflavin-T fluorescence (100 µM compound, 100 µM α-synuclein monomer, n = 3, 24 h). All three engine-nominated candidates inhibited aggregation with a perfect prospective inhibitor-call concordance (3/3), and the EGCG positive control behaved as expected. 2-D08 and EGCG reduced the detectable ThT signal to background levels, Uralenol reduced the 24-h plateau by ~68%, and Herbacetin produced partial inhibition (~51% plateau reduction with a delayed ~12 h lag); two of the four assayed compounds (2-D08 and EGCG) achieved ≥80% plateau reduction (Fig. 3, Table 2). A pilot round at the sub-saturating 1 µM dose showed no statistically significant inhibition, with only weak trends for some compounds (Supplementary Fig. 4); these exploratory data were not used for the prospective concordance analysis and are compatible with a dose-dependent mechanism. Because the goal of this round was prospective validation of the engine rather than lead optimization, we assayed commercially available compounds (no de novo synthesis) for anti-aggregation reactivity only, without imposing the gastrointestinal-absorption or BBB constraints applied elsewhere in the pipeline; among the engine-prioritized commercial chemotypes, 2-D08, Uralenol and Herbacetin were the three our laboratory could actually procure and assay (their ranks within the disclosed commercial panel span 7–22). All four assayed compounds share the engine-predicted primary hit-point at the V66–V77 β-sheet core, with secondary contacts at the G68–A69 hinge and the E83 salt-bridge, mechanistically consistent with the observed phenotypes.

**Table 2.** Engine ↔ wet-lab concordance (α-synuclein ThT, 100 µM, 24 h, n = 3).

| Metric | Result | Interpretation |
| --- | --- | --- |
| Engine-nominated candidates | 3 (+ EGCG ctrl) | prospective calls |
| Observed inhibition (100 $\mu\text{M}$) | 3 / 3 nominated | + EGCG ctrl as expected |
| $\geq 80\%$ plateau reduction | 2 / 4 assayed | 2-D08, EGCG |
| Delayed lag ($\geq 1.5\times$) | 2 / 4 assayed | Herbacetin, EGCG |
| Prospective concordance | 3 / 3 nominated | engine-call validity |

**Figure 3.**
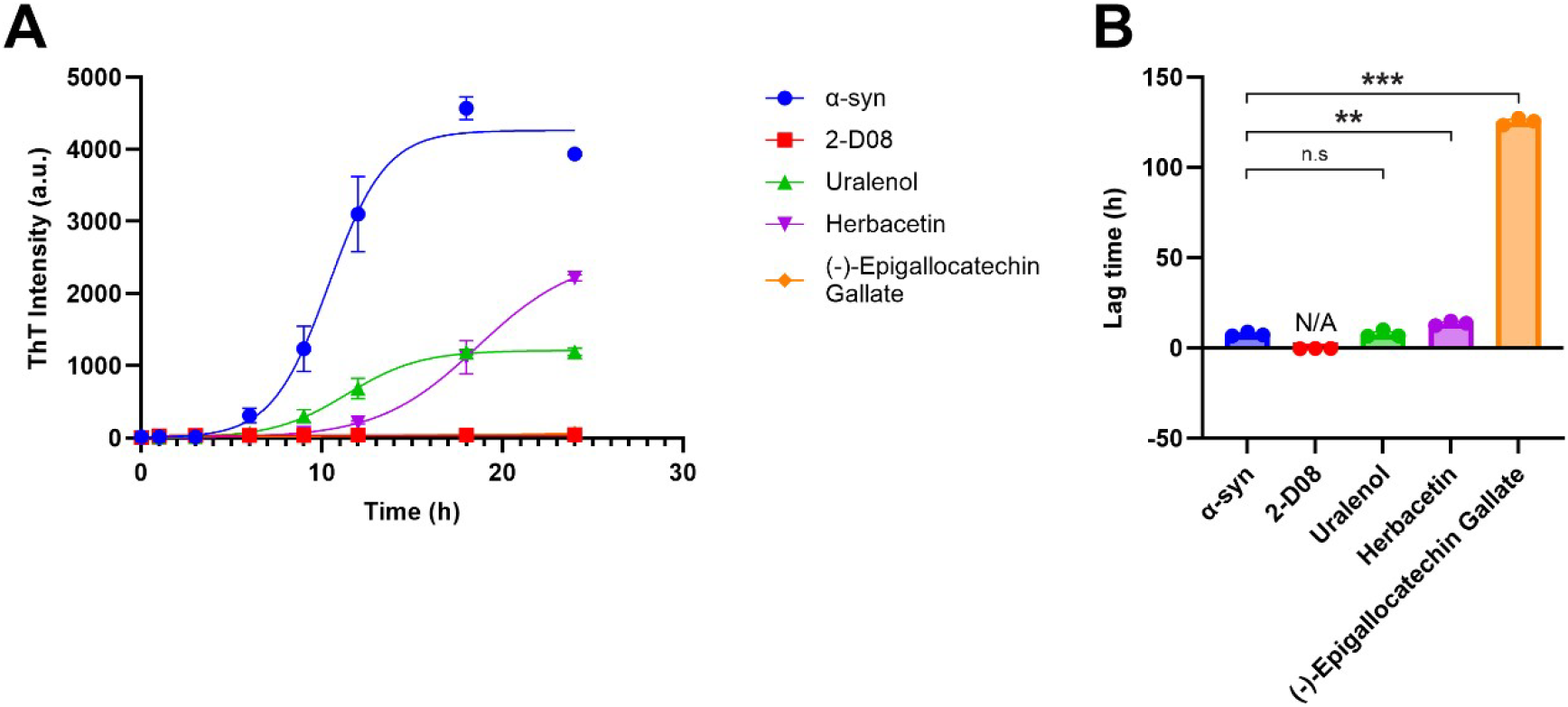
Wet-lab validation by thioflavin-T at 100 µM (mean ± SEM, n = 3). (A) Aggregation kinetics over 24 h: α-synuclein alone (blue) reached a full aggregation plateau, whereas the engine-nominated candidate 2-D08 (red) abolished or strongly suppressed the ThT signal. Uralenol (green) and Herbacetin (purple) partially reduced aggregation (~68% and ~51% inhibition, respectively), while the positive control EGCG (orange) completely abolished aggregation. (B) Apparent aggregation lag times fitted from the kinetic curves: Because 2-D08 abolished aggregation, no lag time could be determined (N/A). Uralenol and Herbacetin both delayed the onset of aggregation, whereas EGCG prevented detectable aggregation within the 24-h window, resulting in an apparent lag time beyond the experimental window. All three engine-nominated candidates inhibited α-synuclein aggregation (3/3); EGCG behaved as the expected positive control. Statistical significance was assessed by one-way ANOVA followed by Bonferroni’s post hoc test for multiple comparison. **P < 0.01; ***P < 0.001; n.s., not significant.

**Figure 4.**
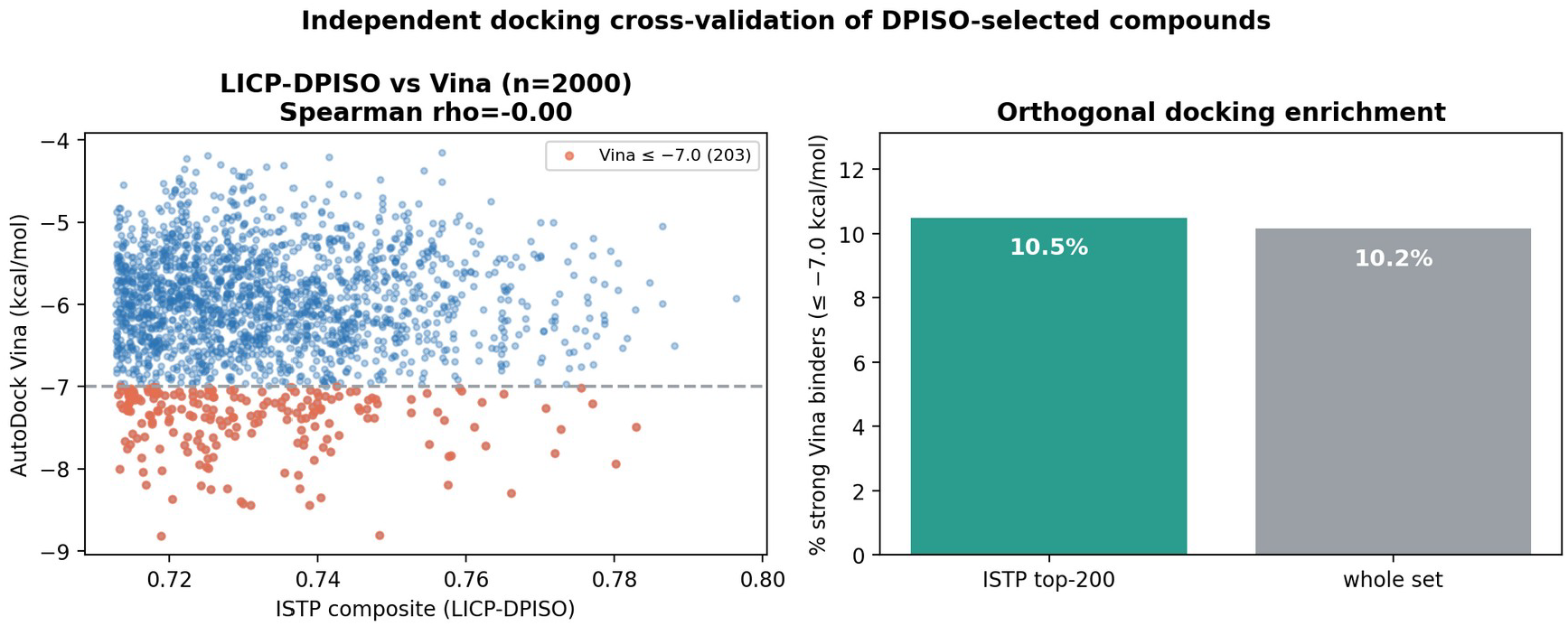
Independent docking cross-validation. Left: ISTP composite (LICP-DPISO) versus AutoDock Vina for 2,000 docked compounds (Spearman rho ~ 0.00). Right: ISTP top-200 shows no docking-based enrichment (10.5% vs 10.2% strong Vina binders), showing no enrichment by rigid-receptor docking in this IDP/fibrilinterface setting and motivating the physics-based score.

### Structural basis of engagement: the NAC hotspots

Engine hotspots were mapped onto the alpha-synuclein NAC fold (PDB 2N0A, residues 61-95; Supplementary Fig. 3). All three ThT-validated compounds (2-D08, Uralenol, Herbacetin) shared a primary contact at the V66-V77 hydrophobic beta-sheet core, with secondary engagement at the G68-A69 nucleation hinge and the acidic E83 salt-bridge. Their chemotypes (flavone/chalcone polyphenols, MW 270-370, cLogP 2.3-3.5, multiple hydroxyls) match this physicochemistry: aromatic rings for pi-stacking in the hydrophobic core and hydroxyl H-bond donors for the E83 electrostatic anchor, which is mechanistically consistent with the observed ThT phenotypes.

### Why DPISO and LICP: rigid-receptor docking is uninformative for this IDP target, and the active set compresses ~ 85x

Classical rigid receptor docking is poorly suited to intrinsically disordered targets with no stable pockets. On a 2,000-compound docked set, the ISTP composite (LICP-DPISO) and AutoDock Vina were essentially uncorrelated (Spearman rho ~ 0.00), and ISTP’s top 200 were no more enriched for Vina strong binders (<= −7.0 kcal/mol) than background (10.5% vs 10.2%; Fig. 4). Therefore, the validity of the engine does not rest on agreement with docking; it rests on calibration (AUC 0.955; known active luteolin/quercetin/baicalein independently recovered) and wet-lab ThT (3/3; nominated + EGCG control). This is exactly the regime in which a physics-based, ensemble-aware score is required; static-receptor docking did not enrich functional aggregation inhibitors in this NAC interface setting, whereas the LICP-DPISO score did. The per-component ablation procedure (-DPISO, -LICP, and -Genesis), together with the qualitative full-engine vs. docking-only contrast, is described in the SI (S4).

#### LICP compression

Across the local ZINC mirror (~8.46 × 10^8^ molecules), the coarse BBB+GI gate plus the LICP capacity-limited active set bound the expensive LICP-DPISO scoring to a 10,000,000-molecule active set -- a ~85x reduction in the number of compounds that reached phase-interference scoring (Supplementary Table 4).

## Discussion

### From hit ranking to state-space-compressed discovery

The central result of this study is not a particular α-synuclein inhibitor but ISTP-DPISO DrugEngine itself: an end-to-end pipeline whose output is already filtered for selectivity, brain penetration, toxicity, synthetic accessibility and chemical novelty, and is closed by wet-lab confirmation. Its defining premise is that the bottleneck in early discovery is not the search algorithm, but the size of the state space that the algorithm must traverse. Rather than dock-ranking a tractable sub-library, the engine first constructs an information-bearing active set and then evaluates candidates in detail so that the scoring effort is spent where it is informative. α-Synuclein is the validation case, not the subject—the contribution of this work is not the discovery of four inhibitors, but the construction of a reusable discovery engine that produced them.

### State-space size is the bottleneck, not scoring

Drug discovery has traditionally focused on improving scoring functions, and the response to ever-larger chemical libraries has been to dock-rank ever-larger sublibraries; ultra-large structure-based campaigns now enumerate and score 10^8^–10^10^ make-on-demand molecules^18–20^. Yet exhaustively scoring a ~8.46×10^8^-molecule mirror with a physics-based, ensemble-aware function is computationally prohibitive on commodity hardware, and rigid-receptor scoring functions are themselves only weakly predictive of true affinity^21^. Our results suggest that the dominant bottleneck is often not scoring accuracy, but the size of the candidate space; a modest improvement in ranking quality cannot compensate for an intractable search space, whereas active-set compression changes the effective problem size itself. The production run makes this concrete — the same composite score that fails to dominate brute force at 8.46×10^8^ molecules becomes tractable once LICP reduces the scored set ~85-fold (Fig. 2). Rather than accelerating the scorer or hierarchically pruning the docking tree^20^, ISTP-DPISO DrugEngine removes molecules that never require scoring; therefore, we frame compression, not scoring, as the primary lever for billion-scale physics-based discovery.

### The DPISO search operator

DPISO is a classical deterministic discrete search operator that is applied after LICP compresses the candidate space into a capacity-limited active manifold. It accumulates phase-like evidence over candidate–hotspot interactions, constructively amplifies coherent low-energy regions, and prunes low-information branches, returning a phase-interference quality that enters the composite score as one term. The computational gain therefore arises primarily from reducing the state space before any high-cost scoring, rather than from accelerating a conventional docking or scoring function: in an astronomically large library, the number of molecules that must be evaluated to find the best candidates is itself the limiting cost, and LICP removes the molecules that never needed scoring before DPISO ranks the remainder.

### LICP compression as the enabling principle

The strongest claim of this study is based on the compression principle. In our previous LICP-ISTP streaming interaction mapping^16^, a high-dimensional interaction structure was captured without materializing a dense N × N matrix. Only block summaries, a magnitude distribution, and the top-K extreme interaction pairs were retained, bounding storage at O(B^2^)+O(K), where dense storage would otherwise grow into the gigabyte range. The same principle is transferred from interaction mapping to chemical screening; instead of materializing the full chemical state space, the engine preserves only the active information-bearing subset. In the production run, this compressed a ~8.46×10^8^-molecule mirror to a 10,000,000-molecule active set (~85-fold) before any expensive scoring, and it is this step, not a faster scorer, that makes billion-scale physics-based screening tractable.

### Connection to compressed-manifold navigation in FeMoco

The same compression thesis was previously demonstrated in a very different setting: active-space reduction for the FeMoco electronic-structure problem, where a formal Hilbert space of dimension ≈ 3.8×10^30^ was navigated with ≈ 7.48×10^5^ determinants, a compression of order 10^24^ — placing the electronic and chemical settings on the same footing^17^. In both electronic and chemical settings, the formal space is misleadingly large, and the useful search space is a structured, low-volume manifold. Whereas FeMoco compresses an electronic configuration space, and the present engine compresses a chemical library space, the operative principle is identical: identifying the small, information-bearing manifold that actually carries the physics and navigates only that, rather than enumerating the formal space. Therefore, ISTP-DPISO DrugEngine extends compressed-manifold navigation from electronic Hilbert spaces to chemical candidate spaces, placing the drug engine, FeMoco work, and our broader LICP-ISTP program within one computational lineage.

### Generality of the compression principle

Because LICP acts on the abstract structure of a candidate space rather than on chemistry, the same principle is expected to apply wherever a formally enormous space hides a structured, low-volume active manifold: protein engineering (sequence–structure space), material discovery (composition–lattice space), quantum chemistry (electronic configuration space, as in FeMoco), and graph or combinatorial optimization. In each case, the recipe is identical: stream the formal space through cheap admissibility gates, compress it to a capacity-limited active set, and search the set with a phase-interference operator. Thus, the drug engine is best read as an instantiation of the general state-space-compression methodology.

### α-Synuclein as a stringent validation target

α-Synuclein was selected precisely because it represents one of the most challenging classes of targets: an intrinsically disordered protein with no stable binding pocket, populating a broad conformational ensemble that even state-of-the-art static structure prediction^22^ represents only as a low-confidence single fold. A method that performs on this target is being tested in the regime where conventional structure-based pipelines are the weakest, which makes prospective 3/3 wet-lab concordance a demanding rather than permissive test. Targets with well-defined pockets should, if anything, be easier for the same engine.

### Why rigid docking fails for α-synuclein

The near-zero correlation between the engine score and AutoDock Vina and the absence of Vina-binder enrichment among the engine’s top 200 (Fig. 4) are not weaknesses but predictable consequences of rigid-receptor docking. AutoDock Vina scores each ligand against a single fixed receptor conformation with a grid-based scoring function and ranks candidates according to the depth of the best static pose^5,6^. This procedure has been well-validated for proteins with a stable, preformed pocket, but its accuracy is known to degrade sharply when the binding site is shallow, flexible or absent^21^. An intrinsically disordered protein such as α-synuclein has no single native fold; it exists as a broad ensemble of rapidly interconverting conformations, and ligands act by subtly shifting that ensemble, rather than docking into a fixed cavity^23^. Consequently, a single static receptor cannot represent the true conformational ensemble, the deepest-static-pose criterion becomes ill-defined, and the docking scores become unstable and only weakly predictive. The engine avoids this failure mode by not treating the target as a fixed object at all: LICP and DPISO operate on a compressed state space of physically accessible interaction configurations rather than on one frozen receptor structure, so the relevant question becomes whether a candidate perturbs aggregation-relevant hotspots while remaining developable and non-reactive — exactly what the engine scores. Therefore, its validity rests on calibration (AUC 0.955; luteolin/quercetin/baicalein independently recovered) and prospective wet-lab confirmation, and not on agreement with docking. More importantly, the engine is not restricted to disordered targets. As an exploratory sanity check on five well-folded drug targets (GSK3β, EGFR, BRAF, MAO-B and HIV-1 protease; 30 known actives and 30 property-matched decoys each, scored by real AutoDock Vina docking and by the engine), the DPISO score gave active/decoy discrimination at least comparable to rigid-receptor docking (Supplementary Fig. 5 and Supplementary Table 5), while on the IDP α-synuclein the engine reached AUC 0.955 where docking was uninformative. This panel is a sanity check rather than a comprehensive docking benchmark; the variable Vina performance is consistent with documented limitations of rigid-receptor docking for virtual-screening enrichment^21^, and indicates that the α-synuclein result reflects target physics rather than an engine artifact.

### Functional validation and calibration limits

All three engine-nominated candidates inhibited aggregation (3/3 at 100 µM; the EGCG positive control behaved as expected), establishing prospective functional validity of the engine call. The ThT result should be read as prospective functional validation, not as final medicinal-chemistry optimization: there is as yet no dose–response IC50, no orthogonal assay (TR-FRET, TEM), the readout is a single, deliberately saturating 100 µM concentration (chosen for an unambiguous kinetic readout against the highly aggregation-prone α-synuclein monomer), and ThT fluorescence interference cannot be fully excluded; dose-response (IC50) titrations, a compound-only ThT interference control and TEM fibril imaging are underway. The quantitative ranking is only partially calibrated—the engine placed Uralenol ≈ Herbacetin above the chalcone/catechin pair, whereas observed potency was 2-D08 ≈ EGCG > Uralenol > Herbacetin—indicating the cross-reactivity penalty for promiscuous polyphenols is under-weighted. The four assayed positives were seeded as hard positives for the next calibration cycle. The uniform HIGH (200/200) cross-reactivity flag is not a failure but a faithful signal of NAC groove non-specificity and is precisely what motivates a de novo first route to purpose-built, selective chemotypes. Notably, the engine’s top-ranked set includes non-polyphenolic, CNS-druggable scaffolds that satisfy both oral-absorption and blood–brain-barrier criteria (Supplementary Information S7), directly addressing the long-standing brain-exposure limitation of the polyphenolic chemotypes that dominate known α-synuclein aggregation inhibitors.

### Implications for prion-like and aggregation-driven targets

Because the engine is configuration driven and unmodified between targets, the same compression plus DPISO procedure is applied to any aggregation-driven or intrinsically disordered target. Beyond polyphenols, structurally unrelated agents such as ceftriaxone also perturb α-synuclein polymerisation^24^, suggesting the hotspot-driven score could generalize across chemotype classes. The next steps are dose–response and orthogonal validation of the present hits, a DPISO/LICP ablation that quantifies each component’s contribution, prospective validation across the engine’s other configured targets, a direct-destabilizer track for prion-like aggregates, and full structural disclosure of the withheld engine hits once patent protection is secured. The broader implication is that the discovery of these targets is rate-limited by state-space size rather than scoring speed, and that an explicit compression step is the decisive lever.

## Methods

### Target preparation

Human α-synuclein coordinates were taken from PDB 2N0A (solid-state NMR fibril, 10 chains, 20,160 atoms) ^10^. The receptor was protonated at pH 7.4 and converted to PDBQT using Open Babel 3.1.0^25^. The binding site was defined at the Chain A–B NAC interface (residues 61–95), grid center [109.5, 130.6, −25.5] Å, 30 × 30 × 30 Å box.

### Stage 1 — Protein Genesis hotspots

The Protein Genesis residue-graph module identifies aggregation hotspots from the NAC sequence; for α-synuclein these are the V66–V77 β-sheet core (primary), the G68–A69 nucleation hinge, and the E83 salt-bridge anchor.

### Stage 2 — DPISO search

Candidate conformational manifolds were traversed by the Discrete Phase-Interference Search Operator (DPISO) on a 128-node grid (300 steps, absorption 0.15), which is a classical, deterministic discrete operator that accumulates phase-like evidence across candidate–hotspot interactions, amplifies coherent low-energy regions, and prunes low-information branches. The resulting phase interference quality was entered into the composite score as a single term.

### Stage 3 — LICP stability gate

The Local Information Criticality Principle (LICP) gate retains candidates whose interaction manifolds are stable under a capacity-limited active-set criterion, and unstable manifolds are rejected.

### Stage 4 — De novo generation

New chemotypes were assembled by directional fragment growth (safe_combine) with two LICP-derived pruning rules: dead-branch termination (abandoning a branch when a child scores < 80% of its parent) and scaffold leaf-peeling^26^ (prune scaffold families with a single analog; calibration scaffolds protected).

### Stage 5 — Cross-reactivity rejection

Each candidate was scored against a 242-hit-point anti-target panel built from 100 seed proteins; a per-compound selectivity gap (target pocket-fit minus maximum off-target pocket-fit) and a HIGH/MED/LOW risk call were emitted. The comparisons used matched pocket-fit scales (never combined vs. pocket-fit).

### Stage 6 — BBB / ADMET

A toxicophore filter applies 38 hard-kill and 7 soft-monitor SMARTS rules (pan-assay interference compounds (PAINS), reactive groups, human ether-à-go-go-related gene (hERG)/cytochrome P450 (CYP) liabilities) ^27^; the ADMET penalty is 0.03 per soft flag (cap 0.15). The BBB-tight CNS configuration additionally requires BBB+ prediction^28^, MW 180–450 Da and cLogP ≤ 4.5^29^. For central-nervous-system targets, candidates are additionally prioritised by a two-tier CNS-druggability filter that combines an oral-absorption tier (Veber/Lipinski) with a blood–brain-barrier CNS-MPO desirability; this metric supersedes the conservative soft BBB flag and is detailed in Supplementary Information S7.

### Stage 7 — Commercial availability

Survivors are cross-referenced against in-stock ZINC tranches^30,31^ and annotated with vendor, catalog number, and price. The commercial arm screens the purchasable subset directly.

### Stage 8 — ZINC reverse novelty

Morgan/ECFP4 fingerprints^32^ (radius 2, 2048 bits, RDKit^33^) were computed for each candidate, and the maximum Tanimoto similarity to the nearest ZINC compound was recorded; < 0.40 denotes a novel chemotype.

### Composite scoring

In IDP mode the composite score is 0.30^·^genesis-landscape + 0.20^·^persistence + 0.20^·^DPISO + 0.15^·^LICP + 0.10^·^QED^34^ + 0.05^·^synthesis^35^, minus the ADMET penalty, multiplied by the gate term. The docking cross-checks used AutoDock Vina 1.2.7^5,6^ (exhaustiveness 8, 5 poses).

### Algorithm and engine interface (reproducibility)

#### Pipeline (Algorithm 1)

Given library L, target hotspot H, and active-set capacity C_active = 10^7^, the pipeline proceeds as follows: First, a coarse gate retains molecules satisfying MW 180–450 Da, cLogP 0–4.5, TPSA at most 90 Å^2^, HBD at most 3, HBA at most 7, and rotatable bonds at most 8^36^. Second, LICP_ selects the streams of retained molecules and constructs a capacity-limited active set A whose size does not exceed C_active. Third, each molecule in A was scored using the DPISO_score against H and fused into composite and target-combined scores. Fourth, the ranked candidates were passed through cross-reactivity (242 anti-targets), BBB/ADMET, commercial availability, and ZINC reverse-novelty gates. Finally, the pipeline emits a safety-gated shortlist and de novo design. The run is deterministic, given a fixed random seed.

#### Operators

LICP_select is a capacity-limited information-criticality selector with three stated properties: bounded output (|A| ≤ C_active), monotonicity (a molecule retained at capacity C is retained at any C′ > C) and determinism given the seed. The DPISO_score is a classical deterministic discrete operator on a 128-node manifold (300 steps, absorption 0.15) that accumulates phase-like evidence along excitation-connected transitions with constructive amplification of low-energy regions and destructive pruning elsewhere.

#### Proprietary core (patent pending)

The internal kernels — the information-criticality weighting inside LICP_select, the phase-accumulation and pruning schedule inside DPISO_score, and the calibrated composite weights — are proprietary technologies of ISTP TECH Co., Ltd. and are withheld. They are provided as a black box executable under a license/material-transfer agreement, together with the disclosed Algorithm 1, operator properties, parameters, orchestration, and calibration harness with active/decoy sets (available on reasonable request), which enables output-level reproduction of the reported screening campaign and rankings without exposing the proprietary kernel source code.

### Computing environment and desktop-scale execution

All production-scale screening and figure generation runs reported herein were executed on a single desktop workstation running Windows 11 Pro (25H2). The system has a 13th-generation Intel Core i5-13600K CPU (3.5 GHz), 32 GB DDR5 RAM (4800 MT/s), 5.46 TB of local storage, and an NVIDIA GeForce RTX 4090 GPU (24 GB). The local ZINC mirror was held on a local disk, and the pipeline ran without access to the high-performance computing clusters. The reported screening run was CPU-driven (multi-core); RTX 4090 was available for auxiliary cheminformatics and visualization, but was not required for LICP active-set construction. The coarse developability gate over the full ~8.46×10^8^-molecule mirror completed in ~22 min, followed by parallel-sharded engine scoring of the active set, for an end-to-end production campaign on the order of tens of minutes in the recorded run. Runtime timestamps, shard logs, processed molecule and active-set counts, thread counts, random seeds, and hardware-environment records are included in the reproducibility package (Table 3).

**Table 3.** Desktop workstation and production-run footprint.

| Item | Value |
| --- | --- |
| Operating system | Windows 11 Pro (25H2), 64-bit |
| CPU | Intel Core i5-13600K, 3.5 GHz (13th gen) |
| RAM | 32 GB DDR5, 4800 MT/s |
| GPU | NVIDIA GeForce RTX 4090, 24 GB (auxiliary) |
| Local storage | 5.46 TB |
| HPC / cluster | none |
| Library mirror | $\sim 8.46 \times 10^8$ ZINC molecules, 1,313 tranches |
| Active-set ceiling | 10,000,000 |
| Retained LICP active set | 10,000,000 |
| Passed developability filter | 4,892,716 (48.9%) |
| Coarse-gate wall-clock | ~22 min (full mirror) |
| End-to-end production campaign | ~30 min |

#### Preparation of recombinant α-synuclein monomers

Full-length mouse α-synuclein proteins were expressed in *E*.*coli* BL21(DE3) cells transformed with the pRK172-mSNCA construct. After incubation at 37°C for 16 h, bacterial cells were harvested by centrifugation at 6,000 × g for 10 min. The bacterial pellet was resuspended in high-salt buffer containing 750 mM NaCl, 10 mM Tris (pH 7.6), 1 mM EDTA, and 1 mM phenylmethylsulfonyl fluoride (PMSF). The resuspended pellet was lysed by sonication for 30 min (10-sec pulses on/off) at 37% amplitude, followed by boiling for 15 min and centrifugation at 6,000 × g for 20 min. The mouse recombinant α-synuclein monomers in the resulting supernatant were purified by serial purification steps using Superdex 200 Increase 10/300 G size-exclusion and Hitrap Q Sepharose Fast Flow anion-exchange columns (Cytiva), the α-synuclein monomers were subjected to HiTrap SP HP cation-exchange column (Cytiva) to remove lipopolysaccharides. Before use, the α-synuclein monomer was kept at −80°C. The wet-lab validation thus used full-length mouse α-synuclein, whose NAC aggregation core is highly conserved relative to the human α-synuclein sequence used for computational target definition (PDB 2N0A).

#### Thioflavin-T assay

Recombinant α-synuclein monomers (100 µM) were incubated with each compound (100 µM; 1 µM in the sub-saturating first round) for 24 h with ThT readout (ex 450 nm / em 510 nm) at 0, 1, 3, 6, 9, 12, 18 and 24 h, n = 3^37^. Inhibition was determined from the 24-h plateau and lag time relative to the α-synuclein-alone control.

### Reproducibility

The engine was configuration driven and unmodified across runs. The complete Run A–F package (configs, orchestrator, and Windows/POSIX launchers) regenerated every computational figure; see Data and Code Availability.

## Acknowledgments

The authors thank the developers of NumPy, CuPy, Numba, and the open-source scientific computing community.

## Author contributions

Conceptualization: D.H.K. Methodology: D.H.K. Software: D.H.K. Investigation (computational screening and de novo design): D.H.K. Investigation (wet-lab thioflavin-T experiments): J.P. Validation: D.H.K., J.P., S.K. Formal analysis: D.H.K. Resources (wet-lab): S.K. Data curation: D.H.K. Visualization: D.H.K. Literature and reference curation: G.-O.K. Supervision: D.H.K. (overall); S.K. (wet-lab validation). Project administration: D.H.K. Writing – original draft: D.H.K. Writing – review & editing: D.H.K., S.K., J.P., G.-O.K.

## Data and code availability

All source data underlying the figures and tables, precomputed numerical outputs, validation logs, and publication-quality figures are publicly available on Zenodo (https://doi.org/10.5281/zenodo.20318165). The α-synuclein receptor coordinates (PDB 2N0A) and the ZINC library accession details are described in the Methods; all commercially available, CAS-confirmed validation compounds are fully disclosed in Supplementary Table 2, while the structures (names/SMILES) of the engine’s novel screening hits are withheld pending patent protection. Pipeline orchestration (the Run A–F driver), coarse-gate thresholds, active-set construction, and figure-generation scripts are released for reproducibility. The proprietary ISTP-DPISO DrugEngine — including the core LICP pruning, DPISO frontier navigation, the de novo generation module, and the LICP/DPISO scoring kernel — is patent-pending and is not deposited in any public repository; it is provided only as a black-box executable, which, together with example input/output pairs and step-by-step instructions enabling independent regeneration of all reported results, can be made available to editors and reviewers upon reasonable request under a confidential evaluation agreement.

## Supplementary Materials

Materials and Methods

Supplementary Text

Figs. S1 to S8

