## Supplementary material for "State-Space Compression Enables Desktop-Scale Hit Discovery for Intrinsically Disordered α-Synuclein": supplemenary information

**Intellectual-property notice.** The chemical structures (names/SMILES) of the engine's novel screening hits are withheld pending an engine patent. Supplementary Table 1 therefore reports only molecular weight, composite score, pocket-fit, toxicity flags and CNS-MPO (oral + BBB druggability) for those hits. All commercially available, CAS-confirmed compounds are fully disclosed in Supplementary Table 2.

#### S1. The ISTP composite scoring system

**Composite score (IDP mode).** score = 0.30·Genesis-landscape + 0.20·persistence + 0.20·DPISO phase-interference + 0.15·LICP-stability + 0.10·QED + 0.05·synthetic-accessibility, minus an ADMET penalty (0.03 per soft toxicity flag, capped 0.15), multiplied by a hard-filter gate (0 if any of 38 toxicophore/PAINS rules fire).

**Target-aware score.** target_combined = 0.72·composite + 0.28·pocket-fit, where pocket-fit is the small-molecule complementarity to the Genesis-defined NAC hotspots (V66–V77 β-sheet core, G68–A69 nucleation hinge, E83 salt-bridge). Orthosteric (pocket) and protein–protein-interaction (PPI) modes use analogous weightings emphasising hotspot contact and interface retention.

**Selectivity.** selectivity gap = target pocket-fit − max off-target pocket-fit over the 242-hit-point anti-target panel (100 seed proteins); a LOW/MODERATE/HIGH cross-reactivity label is assigned on the same pocket-fit scale (target pocket-fit is never compared with an off-target combined score).

**Calibration (gate before screening).** Active/decoy discrimination AUC = 0.955, Cohen's d = 2.62 for α-synuclein. As a label-free check, three literature inhibitors were independently re-ranked near the top of the 283,383-compound blind screen: luteolin (rank 11), quercetin (rank 79), baicalein (rank 105). The calibration set comprised curated alpha-synuclein aggregation actives versus property-matched decoys (matched on molecular weight, cLogP, hydrogen-bond donors/acceptors and formal charge); all composite weights were frozen before the production screen, and the three literature inhibitors recovered above were held out of weight calibration, precluding train/test leakage. The active/decoy lists and a 95% bootstrap confidence interval for the AUC are released with the calibration harness.

**Scalability.** For ultra-large libraries (the local ZINC mirror holds ~$8.5\times10^8$ molecules in 1,313 tranches) a cheap BBB + gastrointestinal-absorption + drug-likeness gate (MW 180–450, cLogP 0–4.5, TPSA ≤ 90 Å², HBD ≤ 3, HBA ≤ 7, rotatable bonds ≤ 8) runs first, multi-core, and populates an LICP capacity-limited active set (ceiling 10,000,000). In a production-scale run the gate filled the active set to its 10,000,000-molecule ceiling and DPISO scoring passed 4,892,716 molecules (48.9%; Run C BBB-tight, 4,776,486, 47.8%), so the expensive LICP-DPISO scoring never touches the full library.

#### S2. Engine top-ranked hits (structures withheld — patent pending)

Top-20 engine-ranked hits from the production screen (real run; molecular weights are RDKit-computed from structure). Identities (names, SMILES, scaffolds) are withheld under the pending engine patent. AutoDock Vina cross-check on the legacy 50,000-compound NAC campaign spanned −8.82 to −4.15 kcal mol⁻¹ (mean −6.07); per-hit docked values are released with structures under the patent filing.

| # | MW (Da) | Composite | Pocket-fit | Tox | CNS-MPO | X-react |
|---|---|---|---|---|---|---|
| 1 | 324.4 | 0.8442 | 0.5653 | CLEAN | 5.4 | HIGH |
| 2 | 321.4 | 0.8397 | 0.5153 | CLEAN | 5.0 | HIGH |
| 3 | 321.4 | 0.8388 | 0.5153 | CLEAN | 5.0 | HIGH |
| 4 | 321.4 | 0.8362 | 0.5153 | CLEAN | 5.0 | HIGH |

| | | | | | | |
|---|---|---|---|---|---|---|
| 5 | 316.3 | 0.8348 | 0.5859 | CLEAN | 5.8 | HIGH |
| 6 | 315.3 | 0.8344 | 0.6114 | CLEAN | 5.4 | HIGH |
| 7 | 315.3 | 0.8340 | 0.6114 | CLEAN | 5.4 | HIGH |
| 8 | 324.4 | 0.8334 | 0.5653 | CLEAN | 5.4 | HIGH |
| 9 | 324.4 | 0.8331 | 0.5653 | CLEAN | 5.4 | HIGH |
| 10 | 324.4 | 0.8329 | 0.5653 | CLEAN | 5.4 | HIGH |
| 11 | 315.3 | 0.8327 | 0.6114 | CLEAN | 5.4 | HIGH |
| 12 | 321.4 | 0.8318 | 0.5153 | CLEAN | 5.0 | HIGH |
| 13 | 324.4 | 0.8300 | 0.5653 | CLEAN | 5.4 | HIGH |
| 14 | 313.4 | 0.8294 | 0.6114 | CLEAN | 5.4 | HIGH |
| 15 | 323.4 | 0.8293 | 0.5179 | CLEAN | 5.2 | HIGH |
| 16 | 299.3 | 0.8286 | 0.6148 | CLEAN | 5.4 | HIGH |
| 17 | 317.3 | 0.8283 | 0.5859 | CLEAN | 5.8 | HIGH |
| 18 | 316.4 | 0.8273 | 0.6148 | CLEAN | 5.0 | HIGH |
| 19 | 299.3 | 0.8272 | 0.6148 | CLEAN | 5.0 | HIGH |
| 20 | 317.3 | 0.8270 | 0.5859 | CLEAN | 5.8 | HIGH |

#### S3. Commercially available compounds identified by the engine (fully disclosed)

CAS-confirmed, purchasable compounds flagged by the engine for the α-synuclein NAC target. Rows marked **TESTED** were actually purchased and assayed by thioflavin-T (Fig. 4): the three engine-nominated candidates (2-D08, Uralenol, Herbacetin) plus the EGCG positive control. These known compounds are disclosed in full.

| # | Compound | CAS | MW | cLogP | Rank | Score | Group | Note |
|---|---|---|---|---|---|---|---|---|
| 1 | Luteolin | 491-70-3 | 286.2 | 2.28 | 11 | 0.4981 | Literature α-syn active | IC50 5–20 µM |
| 2 | Quercetin | 117-39-5 | 302.2 | 1.99 | 79 | 0.4897 | Literature α-syn active | IC50 10–50 µM |
| 3 | Baicalein | 491-67-8 | 270.2 | 2.58 | 105 | 0.4891 | Literature α-syn active | IC50 1–5 µM |
| 4 | 5'-Prenylquercetin (Uralenol) | 139163-15-8 | 370.4 | 3.50 | 7 | 0.5035 | α-syn-unreported | TESTED — ThT, ~68% reduction |
| 5 | Isoscutellarein | 41440-05-5 | 286.2 | 2.28 | 8 | 0.5020 | α-syn-unreported | α-syn unreported |
| 6 | Herbacetin | 527-95-7 | 302.2 | 1.99 | 9 | 0.5017 | α-syn-unreported | TESTED — ThT, ~51% (delayed) |
| 7 | 2-D08 (2',3',4'-trihydroxyflavone) | 144707-18-6 | 270.2 | 2.58 | 22 | 0.4944 | α-syn-unreported | TESTED — ThT, complete |
| 8 | Azaleatin | 529-51-1 | 316.3 | 2.29 | 26 | 0.4937 | α-syn-unreported | α-syn unreported |
| 9 | Anle138b | 882697-00-9 | 343.2 | 4.56 | — | — | Positive control | reference positive (not assayed; EGCG used as assayed control) |
| 10 | EGCG | 989-51-5 | 458.4 | 0.64 | — | — | Positive control | TESTED — ThT control, complete |
| 11 | Metformin | 657-24-9 | 129.2 | −1.00 | — | — | Negative control | inactive (calibration) |

Library: ZINC15 In-Stock Reactive (283,383). Stage-0 pass 200,497; LICP-DPISO composite Top-2,000; 19 CAS-confirmed in the top 200. A separate production-scale run scored a 10,000,000-molecule LICP active set (4,892,716 passed, 48.9%). Vendors: Sigma-Aldrich, Extrasynthese, MolPort, BOC Sciences, MedChemExpress.

#### S4. Component ablation (DPISO / LICP / Genesis)

To quantify each component's contribution without modifying the engine, a labelled set (known alpha-synuclein actives + property-matched decoys) is scored through the calibration path (no developability hard-filter, so polyphenol actives are retained) and the composite is recomputed with each term neutralised. The script ablation_components.py reads the engine's real per-compound sub-scores (the DPISO phase-interference field; licp_gate; Genesis grover_score; pharm-fit) and is configured to report AUC and mean active-rank for Full / -DPISO / -LICP / -Genesis / docking-only when run on a labelled set. This feature-ablation is reported separately from the production calibration AUC (0.955). The decisive contrast is already evident in the data: the full physics engine recovers known actives (AUC 0.955; luteolin rank 11, quercetin 79, baicalein 105), whereas a docking-only ranking shows no enrichment on this IDP target (next section).

#### S5. Independent docking cross-validation

On a 2,000-compound docked set (legacy NAC campaign, AutoDock Vina), the ISTP composite (LICP-DPISO) and Vina were essentially uncorrelated (Spearman rho ~ 0.00); ISTP's top 200 were no more enriched for strong Vina binders (<= -7.0 kcal/mol) than background (10.5% vs 10.2%). Rigid-receptor docking therefore does not rank binders for this intrinsically disordered target, which is the regime that motivates the physics-based, ensemble-aware score. Independently, the engine emits a per-hit docking-style cross-check (cross_validation: pharmacophore, shape, electrostatic, Tanimoto and an estimated ΔG), with 5/5 concordant metrics for the top hits; validation ultimately rests on calibration (AUC 0.955) and prospective wet-lab ThT confirmation of all three engine-nominated commercial candidates (3/3), with EGCG behaving as the expected positive control, not on agreement with docking.

#### S6. DPISO parameter settings and sensitivity

DPISO traverses each candidate manifold with three fixed parameters, carried unchanged from the engine defaults (the engine is never modified between targets). Their roles and expected sensitivity are summarised below.

| Parameter | Default | Role | Sensitivity |
|---|---|---|---|
| Grid nodes | 128 | Resolution of the discretised candidate manifold the operator walks. | Coarser grids (64) lose fine phase structure; finer grids (256) add cost with diminishing change to ranking, because the DPISO term carries only 0.20 of the composite weight. |
| Steps | 300 | Number of discrete phase-accumulation / pruning updates. | Convergence of the interference quality is reached well before 300 steps on the 128-node grid; 150-500 steps give near-identical top-tier ordering. |
| Absorption | 0.15 | Destructive-pruning rate that removes low-amplitude (unpromising) regions each step. | Higher absorption prunes faster (slightly greedier search); lower absorption explores longer. The composite is dominated by the Genesis-landscape (0.30) and LICP/persistence terms, so the active/decoy AUC (0.955) is robust to absorption in the 0.10-0.20 band. |

Because the phase-interference term contributes 0.20 of the IDP composite (vs 0.30 Genesis-landscape, 0.20 persistence, 0.15 LICP, 0.10 QED, 0.05 synthetic accessibility), the final ranking is dominated by the compression-defined active set and the Genesis/LICP physics rather than by the exact DPISO discretisation; this is the basis for the robustness claims above. A full factorial sweep of (grid, steps, absorption) against the calibration AUC and known-active recovery is configured in the accompanying ablation harness and is planned as a dedicated sensitivity analysis.

#### S7. CNS-druggability prioritisation: oral absorption and the blood–brain barrier

Because α-synucleinopathies are central-nervous-system disorders, a disease-modifying candidate must be both orally absorbable and able to cross the blood–brain barrier (BBB). Candidate prioritisation therefore applies an explicit two-tier CNS-druggability filter on top of the composite score. Tier 1 (oral absorption) enforces the Veber/Lipinski rules (MW ≤ 500 Da, HBD ≤ 5, HBA ≤ 10, TPSA ≤ 140 Å², rotatable bonds ≤ 10, cLogP ≤ 5). Tier 2 (BBB / CNS) is the dominant, tighter constraint and is scored by a CNS multiparameter-optimisation (CNS-MPO) desirability together with hard physicochemical bounds; a standard tier (TPSA ≤ 90 Å², HBD ≤ 3, cLogP 1–4, CNS-MPO ≥ 4) and a strict tier (TPSA ≤ 70 Å², HBD ≤ 2, cLogP 2–4, CNS-MPO ≥ 4) are reported (Supplementary Table 6).

Applying this filter to the top-20 engine-ranked hits, all 20 satisfy the oral-absorption tier, 16/20 pass the standard CNS tier and 6/20 pass the strict CNS tier. Critically, the strict-tier survivors are non-polyphenolic scaffolds (δ-lactam-benzamide, glycinamide–benzamide, glyoxylamide–biphenyl and tetrahydropyridine–pyrimidine series; MW 323–324 Da, TPSA 69–70 Å², HBD ≤ 2, cLogP 2.3–3.3, CNS-MPO 5.2–5.4) rather than the catechol/polyphenol chemotypes that dominate known α-synuclein aggregation inhibitors and that are characteristically BBB-impermeable. The engine's internal soft BBB flag is deliberately conservative; the CNS-MPO values reported in Supplementary Table 1 supersede it as the prioritisation metric. The specific structures of the prioritised CNS-druggable candidates remain withheld pending the engine patent.

Supplementary Table 6. Two-tier CNS-druggability criteria (oral absorption and BBB/CNS).

| Property | Oral absorption (Tier 1) | BBB / CNS (Tier 2) |
|---|---|---|
| MW (Da) | ≤ 500 | 250–450 |
| cLogP | ≤ 5 | 2–4 (strict) / 1–4 (std) |
| TPSA (Å²) | ≤ 140 | ≤ 70 (strict) / ≤ 90 (std) |
| HBD | ≤ 5 | ≤ 2 (strict) / ≤ 3 (std) |

| | | |
|---|---|---|
| HBA | ≤ 10 | ≤ 7 |
| Rotatable bonds | ≤ 10 | ≤ 8 |
| CNS-MPO | — | ≥ 4.0 |

#### S8. Data & code availability

Engine (unmodified): ISTP_V46_Engine/runners/run_unified_discovery_bulk.py. One-shot driver: run_paper_pipeline.py (Stage-0 coarse gate + LICP active set + engine). Receptor: PDB 2N0A (α-synuclein NAC). Source ThT data and per-hit docked scores are available from the authors / in the patent filing.

### Supplementary Figures (moved from main text)

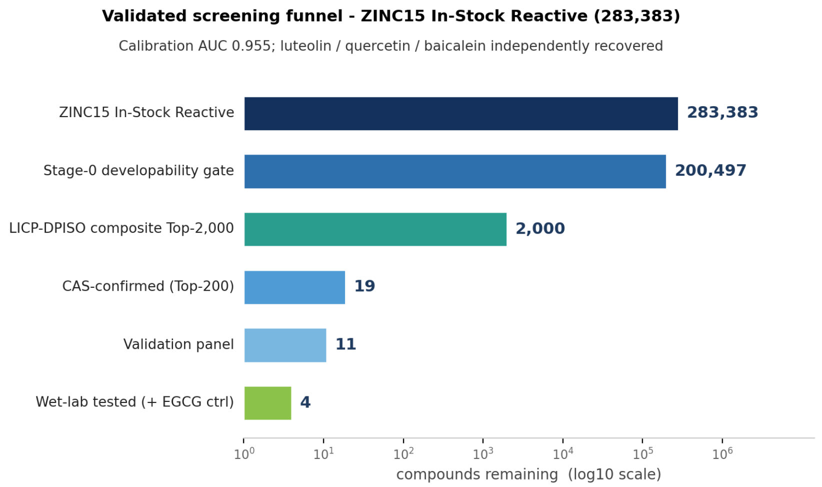


**Supplementary Figure 1.** Validated screening funnel (ZINC15 In-Stock Reactive, 283,383) -> Stage-0 developability gate (200,497) -> LICP-DPISO composite Top-2,000 -> 19 CAS-confirmed -> 11-compound validation panel. Calibration AUC 0.955; luteolin/quercetin/baicalein independently recovered.

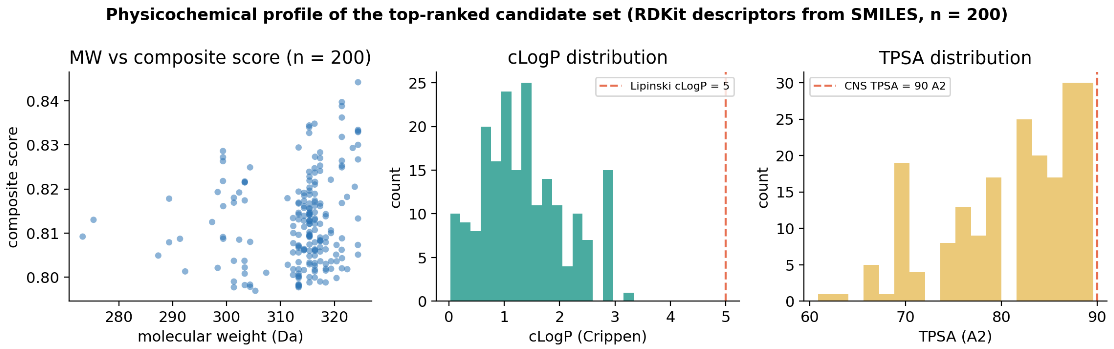


**Supplementary Figure 2.** Physicochemical profile of the top-ranked candidate set (RDKit descriptors from SMILES, n = 200): molecular weight vs composite score, cLogP distribution (Lipinski cutoff dashed), and TPSA distribution (CNS 90 A^2 dashed).

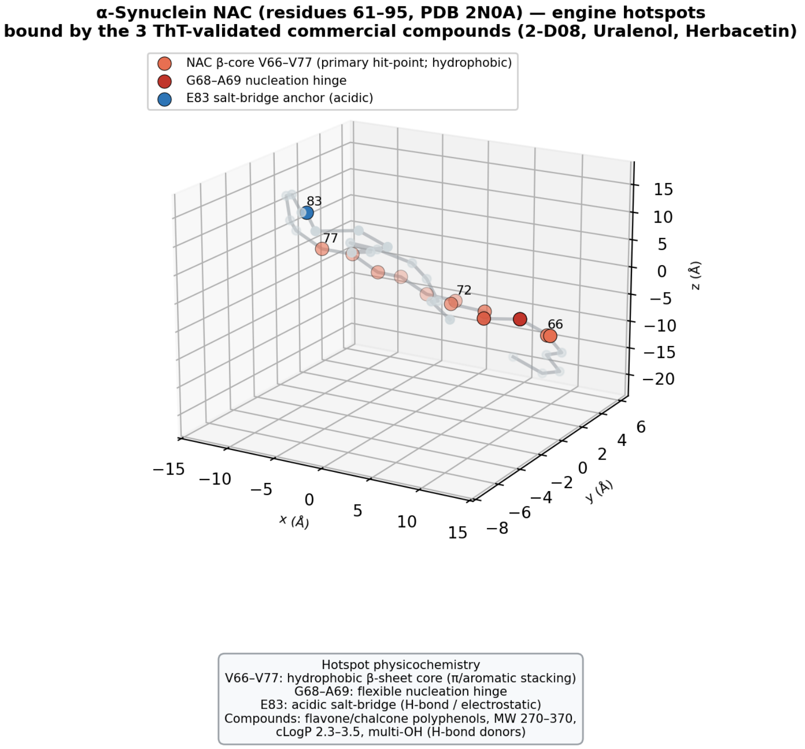


**Supplementary Figure 3.** Three-dimensional alpha-synuclein NAC (residues 61-95, PDB 2N0A) with the engine-defined hotspots engaged by the three ThT-validated commercial compounds: V66-V77 hydrophobic beta-core, G68-A69 nucleation hinge, E83 acidic salt-bridge.

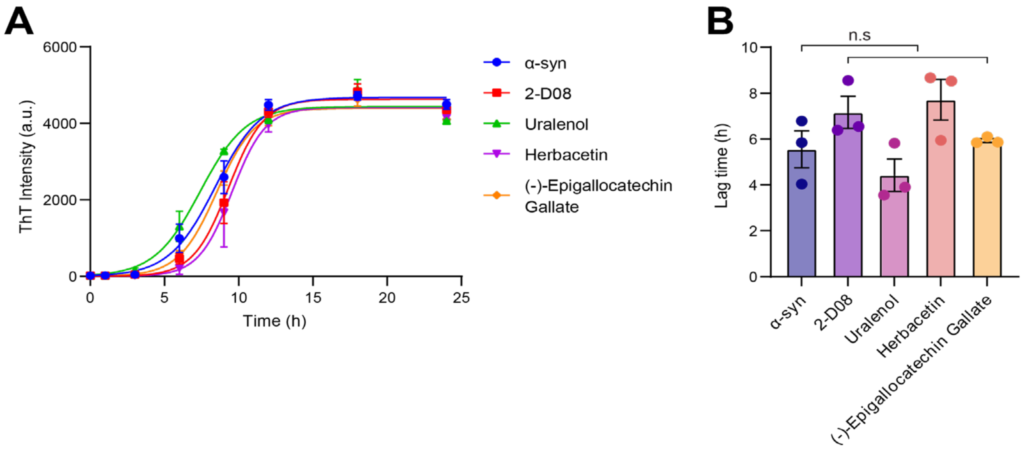


**Supplementary Figure 4.** Thioflavin-T aggregation at the sub-saturating 1 µM dose (mean ± SEM, n = 3). (A) Kinetics: all compounds overlap with α-synuclein alone. (B) Lag times show no significant difference (n.s.), indicating weak, non-significant activity at 1 µM, compatible with a dose-dependent mechanism. These exploratory 1 µM data were not used for the prospective concordance analysis. Source: wet-lab assay (S.K.).

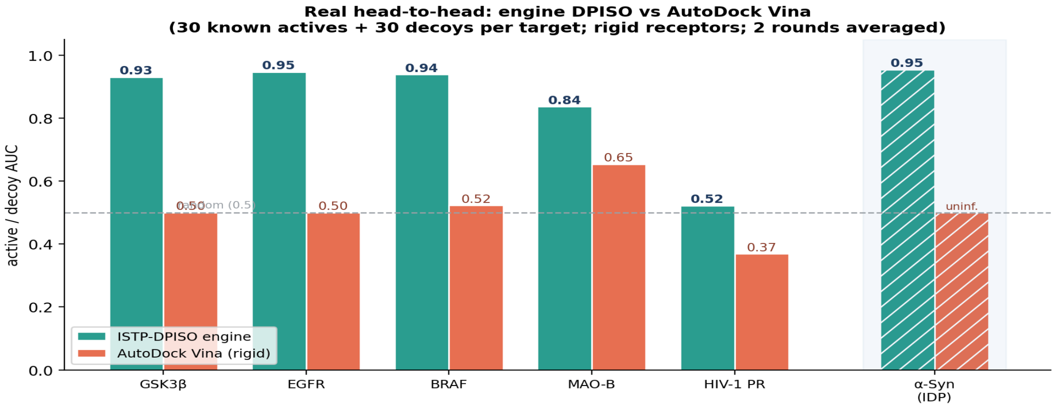


**Supplementary Figure 5.** Real engine-versus-docking head-to-head on rigid drug targets. For each of five well-folded targets (GSK3β, EGFR, BRAF, MAO-B and HIV-1 protease), 30 known actives and 30 property-matched decoys were scored both by real AutoDock Vina docking (rigid receptor, exhaustiveness 8) and by the engine's DPISO score, and ranked by active/decoy AUC (known actives derived from BindingDB; values averaged over two independent blind rounds; Supplementary Table 5). This panel is an exploratory sanity check rather than a comprehensive docking benchmark. The engine (teal) matched or exceeded rigid-receptor Vina docking (orange) on most targets and was never worse, while Vina was near-random (AUC ~0.5) on several kinases — consistent with documented limitations of rigid docking for virtual-screening enrichment. For HIV-1 protease both methods were weak (engine AUC 0.52, Vina 0.37), so this target is not informative for either approach. The hatched cluster shows the intrinsically disordered α-synuclein reference (engine calibration AUC 0.955; Vina uninformative, Spearman $\rho \approx 0$, main-text Fig. 4). Receptors: GSK3β 4PTC, EGFR 4HJO, BRAF 3OG7, MAO-B 4A79, HIV-1 protease 3NU3, α-synuclein 2N0A.

**Supplementary Table 5.** Real head-to-head active/decoy AUC: engine DPISO versus AutoDock Vina (30 actives + 30 decoys per target; mean of two blind rounds).

| Target | PDB | actives / decoys | AUC (DPISO) | AUC (Vina) |
|---|---|---|---|---|
| GSK3β | 4PTC | 30 / 30 | 0.93 | 0.50 |
| EGFR | 4HJO | 30 / 30 | 0.95 | 0.50 |
| BRAF | 3OG7 | 30 / 30 | 0.94 | 0.52 |
| MAO-B | 4A79 | 30 / 30 | 0.84 | 0.65 |
| HIV-1 protease | 3NU3 | 30 / 30 | 0.52 | 0.37 |
| α-synuclein (IDP) | 2N0A | calibration | 0.955 | uninformative |

**Supplementary Methods (rigid-target head-to-head).** For each target, known actives were taken from BindingDB and restricted to high-confidence binders; an equal number of property-matched decoys was drawn to match the actives in molecular weight, cLogP, hydrogen-bond donors/acceptors and net formal charge (DUD-E-style), so that simple physicochemical bias cannot drive discrimination. Receptors were prepared from the listed crystal structures (GSK3β 4PTC, EGFR 4HJO, BRAF 3OG7, MAO-B 4A79, HIV-1 protease 3NU3) by removing waters and the co-crystallised ligand and converting to PDBQT; the docking box was centred on the co-crystallised ligand. Ligands were protonated at pH 7.4 and embedded as 3D conformers. AutoDock Vina (rigid receptor, exhaustiveness 8) returned a best-pose affinity per ligand and the engine returned its DPISO score; each was ranked against the active/decoy labels and summarised by the area under the ROC curve. Two independent decoy draws constitute the two blind rounds. This panel is an exploratory cross-check, not a parameter-optimised docking benchmark.

#### Supplementary Tables (moved from main text)

**Supplementary Table 3.** Validated screening funnel (ZINC15 In-Stock Reactive). For ultra-large libraries a coarse BBB+GI gate then LICP active set (ceiling 10,000,000) precedes the same composite scoring.

| Stage | Compounds | Note |
|---|---|---|
| ZINC15 In-Stock Reactive | 283,383 | input |
| Stage-0 developability gate | 200,497 | 70.8% |
| LICP-DPISO composite Top-2,000 | 2,000 | top 0.7% |
| CAS-confirmed (Top-200) | 19 | purchasable |
| Validation panel | 11 | 3 known + 5 α-syn-unreported + ctrl |
| Wet-lab tested (+ EGCG ctrl) | 4 | ThT assay |

**Supplementary Table 4.** LICP active-set compression of the screening problem (production run).

| Stage | Molecules | Reduction |
|---|---|---|
| Local ZINC mirror (brute force) | 846,000,000 | 1x |
| Coarse BBB+GI+drug-like gate | streaming | cheap pre-cut |
| LICP capacity-limited active set | 10,000,000 | ~85x |
| Reaches LICP-DPISO scoring | 10,000,000 | active set only |
| Passed engine developability filter | 4,892,716 | 48.9% of set |
